# DNA-validated parthenogenesis: first case in a captive female Cuban boa (*Chilabothrus angulifer*)

**DOI:** 10.1101/751529

**Authors:** Fernanda Seixas, Francisco Morinha, Claudia Luis, Nuno Alvura, Maria dos Anjos Pires

**Affiliations:** Laboratório de Histologia e Anatomia Patológica, Escola de Ciências Agrárias e Veterinárias, CECAV- University of Trás-os-Montes and Alto Douro, Quinta de Prados, 5000-801 Vila Real, Portugal; Department of Evolutionary Ecology, National Museum of Natural Sciences (MNCN), Spanish National Research Council (CSIC), José Gutiérrez Abascal 2, 28006 Madrid, Spain; Zoo da Maia, Rua da Igreja, S/N, 4470-184 Maia, Portugal

**Author notes:** Corresponding author. (MAP).

**Keywords:** parthenogenesis, snake, DNA, reptile, *Chilabothrus angulifer*

## Abstract

Parthenogenesis is a biological process of asexual reproduction. Recent studies have highlighted the significance of this fascinating phenomenon in the vertebrate evolution. Although parthenogenetic reproduction appears to be widespread among reptiles, a restricted number of cases were reported in captivity and wild. Here, we studied and reported an intriguing case of a 20-year old captive female Cuban boa (*Chilabothrus angulifer*), from the Zoo da Maia (Maia, Portugal) collection, isolated from conspecifics males, that gave birth twice in 4 years. The neonates from both deliveries, one fresh and the other fixed in formalin, were submitted to histopathological and molecular genetic analysis. Both neonates were homozygous for the loci analyzed, carrying only mother alleles. Furthermore, morphological abnormalities (anophthalmia) were observed in the second neonate. Our data support a pattern of parthenogenetic reproduction. This is the first documented case of facultative parthenogenesis in a Cuban boa, which can be of great interest for further research on ecology, evolution, captive breeding and conservation of the species.

## Introduction

Parthenogenesis is a natural form of asexual reproduction in which offspring is produced from unfertilized eggs [1,2]. This uncommon reproductive strategy was reported in less than 0.1% of vertebrate species, including a wide range of taxa (i.e. fishes, amphibians, reptiles, birds and mammals), even in wild populations [1–5]. Parthenogenetic reproductive events have caught the attention of evolutionary and conservation biologists, since the absence of genetic recombination accelerate the accumulation of deleterious mutations in parthenogenetic individuals, which have considerable implications for the management and conservation of the species [6–8].

Obligate parthenogenesis is a biological process where all individuals within a species reproduce asexually [2,9]. This reproductive strategy is restricted to Squamate reptiles, being reported in various lizards’ species and one snake (*Indotyphlops braminus*) [9–11]. The occasional occurrence of parthenogenesis in individuals of a species that normally reproduce sexually (i.e. facultative parthenogenesis - FP) was first mentioned in the late 1800s for birds [12]. FP have been reported in various species of major vertebrate groups including reptiles, birds and elasmobranchs (sharks and rays) [4,9,13,14]. Most FP events were documented from captive females after long periods without contact to male conspecifics during their reproductive lifetime [8,9]. However, parthenogenesis has more recently been reported in wild snake populations [3,15] and females housed with males [16], suggesting that its occurrence may be more frequent than previously thought in vertebrates. In addition, the reproductive viability of parthenogenetic offspring was observed in some species, which highlights the ecological and evolutionary significance of this reproductive strategy [15]. Nevertheless, the biological basis and mechanisms underlying parthenogenesis remain mostly unknown [4,9,17].

FP was described in at least six snake families, namely Boidae, Pythonidae, Viperidae, Acrochordidae, Colubridae and Elapidae [5,9,17]. Among boid snakes (Boidae family), parthenogenesis was recorded and validated by genetic analysis in *Boa constrictor* and *Eunectes murinus*, as well as, in two species of the genus *Epicrates* (*Epicrates maurus* and *Epicrates cenchria*) closely related to the genus *Chilabothrus* [17–20]. FP confirmed by genetic analysis was also reported for Pythonidae (*Python bivittatus, Python regius* and *Malayopython reticulatus*) [16,21], Viperidae (*Agkistrodon contortrix*) [15,22], Colubridae (*Thamnophis marcianus* and *Thamnophis couchii*) [23,24] and Elapidae (*Oxyuranus scutellatus* and *Acanthophis antarcticus*) [5]. The accurate identification and characterization of parthenogenesis in captive individuals of non-model species may provide important data to understand the frequency, causes, consequences and biological mechanisms of asexual reproduction among vertebrates [1,25].

Two interesting reproductive events were recorded for a captive female Cuban boa (*Chilabothrus angulifer*) isolated from males for eleven years. These occurrences could be explained by two hypotheses: (i) long-term sperm storage from the last mating or (ii) parthenogenetic reproduction. Here, we applied molecular and histopathological methodologies to evaluate these hypotheses, providing the first evidences of facultative parthenogenesis in a Cuban boa.

## Material and methods

### Specimen history and sampling

On 20 September 2017 a 20-year old captive female Cuban boa (*Chilabothrus angulifer* or *Epicrates angulifer*) from the Zoo da Maia (Maia, Portugal) collection gave birth to a stillborn and multiple non-embryonated eggs. This female, purchased to the zoological collection on 1999, had no contact with a male since 2006, when the conspecific male died. Previously, in 2013, this same female delivered a yellowish mass of non-developing eggs and a dead neonate that has been preserved on 10% buffered formalin. The offspring from these two deliveries, one fresh and another formalin fixed, were analysed in the Histology and Anatomical Pathology Laboratory of Trás-os-Montes e Alto Douro University (UTAD). Tissues samples were processed for histopathology according routine technique for light microscopy and staining with haematoxylin and eosin (HE).

### DNA extraction and microsatellite genotyping

The DNA isolation from the formalin-fixed specimen (neonate 2013) was carried out using the Quick-DNA Miniprep Plus Kit (Zymo Research) according to manufacturer’s protocol, with some additional steps before sample digestion. Briefly, a mixture of different tissues (liver, lung, gut and skin) was sliced into small pieces with a scalpel. Then, the tissues were washed with PBS during 24 h (the buffer was replaced twice). The DNA extraction from muscle tissues of the neonate borne at 2017 was performed using the NZY Tissue gDNA Isolation kit (Nzytech). The mother’s DNA was isolated from blood using the NZY Blood gDNA Isolation kit (Nzytech). Both extractions were performed following the standard protocols recommended by the manufacturer.

Thirteen microsatellite markers previously characterized for boid species were analysed: μsat 1, μsat 10, μsat 13, μsat 24, μsat 32, μsat 36, Ci25, Ci34, Ci35, Ci36, Ci37, 55HDZ554 and 55HDZ617 [19,26–28]. The pre-screening of microsatellite variations among mother and offspring samples were performed using high-resolution melting (HRM) analysis [29,30]. PCR amplification and melting acquisition were carried out using a QuantStudio 3 Real-Time PCR System (Applied Biosystems). The reaction mixture was prepared in a 20 μl final volume containing 10 μl of MeltDoctor HRM Master Mix (Applied Biosystems), 5 pmol of each primer and 5 ng of genomic DNA. All PCR reactions were performed in duplicate.

The amplification protocol was run as follows: 1 cycle of 95 °C for 10 min; 40 cycles of 95 °C for 15 s, 60 °C for 1 min (fluorescence signal was captured at the end of each cycle); 1 cycle of 95 °C for 15 s, 60 °C for 1 min and then sequential temperature increments of 0.025 °C/s with temperature ranging from 60 °C to 95 °C, with continuous fluorescence measurements. The melting curve data were analysed with the QuantStudio Design & Analysis software v.1.4 (Applied Biosystems) and High Resolution Melting (HRM) Software v.3.0.1 (Thermo Fisher Scientific), assessing differences in melting curve shapes to characterize microsatellite allelic variability. Forward primers of the microsatellites with variations among samples were labelled with 6-FAM to determine the genotypes using capillary electrophoresis. PCR amplifications were performed in a total volume of 20 μl containing 10 μl of 2x MyTaq HS Mix (Bioline), 5 pmol of each primer and 5 ng DNA. PCR thermal conditions were as follows: initial denaturation at 95 °C for 5 min, followed by 35 cycles of 95 °C for 30 s, 60 °C for 1 min, 72 °C for 30 s and a final extension at 60 °C for 10 min. Amplified fragments were electrophoresed on an ABI PRISM 3130xl Genetic Analyzer (Applied Biosystems) using the GeneScan 500 LIZ size standard. Allele sizes were determined using Peak Scanner v.3.0.2 (Thermo Fisher Cloud).

## Results

Macro and microscopically, both reptiles correspond to fully develop stillborn snakes that showed no morphological alterations except in the 2017 specimen (#2) that present bilateral anophthalmia (Fig 1). Microscopic examination of organs showed no alterations and the presence of the reproductive system confirmed both to be female.

**Fig 1.**
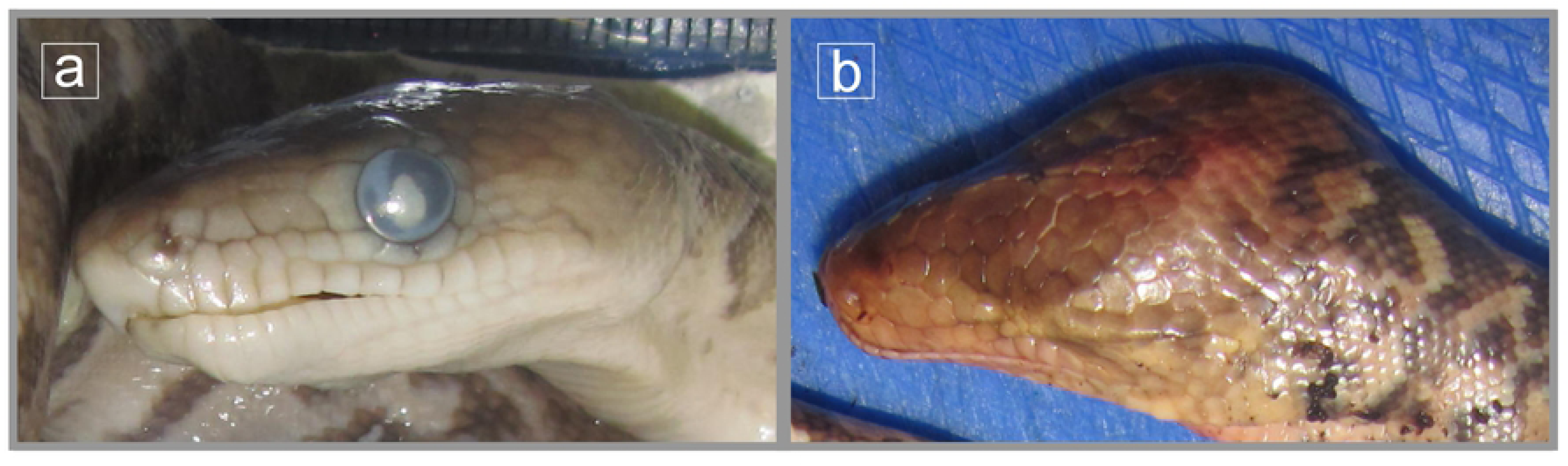
Neonates of Cuban boa (*Chilabothrus angulifer*): (a) Neonate of 2013 with a normal head; (b) Neonate of 2017 with bilateral anophthalmia.

Of the 13 microsatellite loci screened, two markers (μsat 13 and Ci34) did not amplify or generated non-specific PCR products. The high-resolution melting (HRM) analysis allowed the identification of four microsatellite loci (μsat 10, μsat 24, Ci36 and Ci37) with allelic variability among mother and offspring samples (Fig 2). No evidence of allelic variability was detected in the remaining loci for the samples analysed (Fig 2). These results were validated using capillary electrophoresis to determine allele sizes for all polymorphic markers and two non-polymorphic loci (Table 1). Maternal heterozygosity was observed for polymorphic loci and a homozygosity pattern was obtained for the non-polymorphic loci analysed (Table 1). The offspring was homozygous for all microsatellite loci, always carrying an allele present in the mother (Table 1). The locus Ci37 is a potential null allele in the neonate of 2013, since the amplification failed using different DNA samples and PCR conditions.

**Fig 2.**
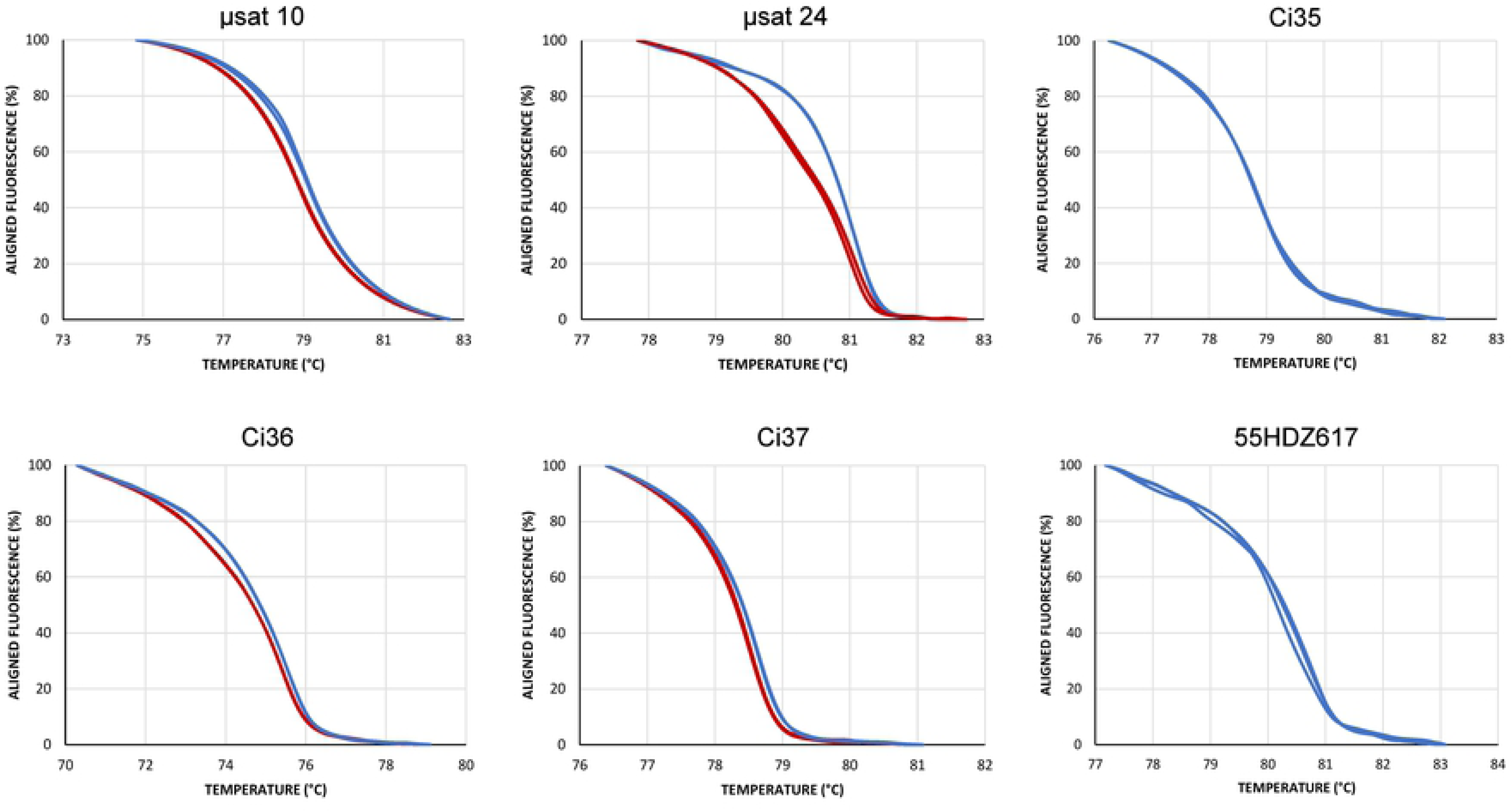
Melting curve profiles obtained in the pre-screening of the microsatellite loci using HRM analysis. The fluorescence differences in four loci (μsat 10, μsat 24, Ci36 and Ci37) allowed the accurate differentiation of the mother and offspring genotypes (red and blue curves). No significant fluorescence variations were detected in loci with same genotype in mother and offspring (Ci35 and 55HDZ617).

**Table 1.**
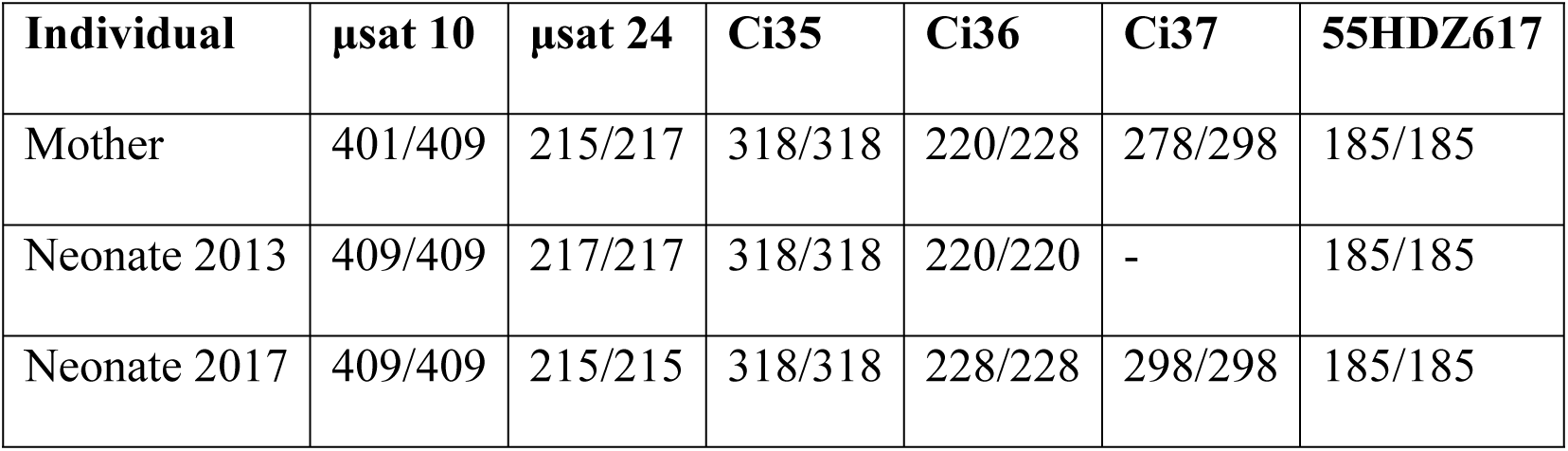
Genotypes of the mother and offspring characterized for the potentially parthenogenic Cuban boa (*Chilabothrus angulifer*)

| Individual | μsat 10 | μsat 24 | Ci35 | Ci36 | Ci37 | 55HDZ617 |
| --- | --- | --- | --- | --- | --- | --- |
| Mother | 401/409 | 215/217 | 318/318 | 220/228 | 278/298 | 185/185 |
| Neonate 2013 | 409/409 | 217/217 | 318/318 | 220/220 | - | 185/185 |
| Neonate 2017 | 409/409 | 215/215 | 318/318 | 228/228 | 298/298 | 185/185 |

## Discussion

Our results support the first evidence of FP in the Cuban boa (*Chilabothrus angulifer*). The allele homozygosity in the offspring, which carries only alleles present in the mother, are similar to previous cases described of FP reported for boid species [17–20]. In Boidae family (for genera Boa and Epicrates), the parthenogenesis was initially detected in all-female litters [18,19]. Similarly, the two stillborn analysed in this study were both females. Accidental FP in captive individuals generally occurs after long periods of isolation from mates [9]. The female in this study had no contact with a conspecific male for 13 years now. Prolonged sperm storage has been documented in various snake species, but the longest time period of suspected sperm storage reported for a snake was seven years and six months [31–33]. In this case the molecular analysis demonstrated the lack of male genetic contribution to the offspring excluding prolonged sperm storage. The high levels of homozygosity detected in the offspring is a characteristic of the parthenogenetic mode explained by terminal fusion automixis [18,19] as recently inferred for long-term captive copperhead (*Agkistrodon contortrix*) and cottonmouth (*Agkistrodon piscivorus*) [5,17].

The genome wide homozygosity of parthenogenetic offspring may be related to the development of malformations [3]. Embryos and stillborn offspring with developmental abnormalities (e.g. anophthalmia, microphthalmia, encephalocoele and head foreshortening) has been associated with parthenogenetic events in reptiles [5,9,34]. The morphological evaluation of the Cuban boa neonate born in 2017 evidenced bilateral anophthalmia, a malformation found in parthenogenetic offspring of other reptile species [34].

In conclusion, we characterized the first record of FP in the Cuban boa supported by specimen history, histological analysis and molecular markers. This may have important ecological and evolutionary implications, being interesting to understand the frequency of this reproductive strategy in captivity, and maybe in the wild, as recorded for some species [3,15,35]. The increasing number of reports on parthenogenic births in a wide range of snakes and other vertebrates also evidences the evolutionary significance of this reproductive phenomena still poorly understood, being a research field with high potential [8,36]. Therefore, we incentive zoo workers, veterinarians, curators, wildlife managers and researchers to pay attention to evidences of abnormal births in this species and related taxa, since these events can be easily neglected.

## Acknowledgements

F.M. was supported by a Juan de la Cierva postdoctoral fellowship (FJCI-2017-32055).

This work was funded by the project UID/CVT/00772/2019 supported by the Portuguese Science and Technology Foundation (FCT).

## Conflict of Interest Statement

The authors declare that the research was conducted in the absence of any commercial or financial relationships that could be construed as a potential conflict of interest.

## Author Contributions

All authors listed have made a substantial, direct and intellectual contribution to the work, and approved it for publication:

Fernanda Seixas: Conceptualization, Investigation, Validation, Writing – Original Draft Preparation

Francisco Morinha: Formal Analysis, Investigation, Validation, Writing – Original Draft Preparation

Claudia Luis, Nuno Alvura: Resources, Writing – Original Draft Preparation

Maria dos Anjos Pires: Funding Acquisition, Supervision, Writing – Review & Editing

